# Keratins determine network stress responsiveness in reconstituted actin-keratin filament systems

**DOI:** 10.1101/2020.12.27.424392

**Authors:** Iman Elbalasy, Paul Mollenkopf, Cary Tutmarc, Harald Herrmann, Jörg Schnauß

## Abstract

The cytoskeleton is a major determinant of cell mechanics, a property that is altered during many pathological situations. To understand these alterations, it is essential to investigate the interplay between the main filament systems of the cytoskeleton in the form of composite networks. Here, we investigate the role of keratin intermediate filaments (IFs) in network strength by studying *in vitro* reconstituted actin and keratin 8/18 composite networks via bulk shear rheology. We co-polymerized these structural proteins in varying ratios and recorded how their relative content affects the overall mechanical response of the various composites. For relatively small deformations, we found that all composites exhibited an intermediate linear viscoelastic behavior compared to that of the pure networks. In stark contrast, the composites displayed increasing strain stiffening behavior as a result of increased keratin content when larger deformations were imposed. This strain stiffening behavior is fundamentally different from behavior encountered with vimentin IF as a composite network partner for actin. Our results provide new insights into the mechanical interplay between actin and keratin in which keratin provides reinforcement to actin. This interplay may contribute to the overall integrity of cells, providing an explanation for the stability of stressed epithelial tissues due to their high keratin contents. Additionally, this helps us to understand the physiological necessity to exchange IF systems during epithelial-mesenchymal transition (EMT) in order to suppress strain stiffening of the network, making cells more elastic and, thus, facilitating their migration through dense tissues.

## Introduction

The complex mechanical behavior of eukaryotic cells is largely determined by the three principal filament systems of the cytoskeleton, i.e. actin filaments (F-actin), microtubules and intermediate filaments (IFs) as well as their regulated interplay.^1^ In addition to establishing a cell’s specific functional and tissue-related shape, these components and their distinct structures are also crucial for numerous general cellular processes such as stability within tissues as well as cell motility, division, and signal transduction.^2^ Alterations of these key constituents are the cause of numerous human diseases. It is somehow perplexing that these very same cytoskeletal components can lead to very static, stable cell conformations, which allow, for instance, to form stable tissue layers such as the epithelium, as well as highly dynamic cells, for instance, during wound healing and cancer metastasis.^3^ Switching between these physically seemingly contradictory cell states highly depends on the ratios of the individual cytoskeletal components. This becomes especially apparent during embryogenesis when cells need to constantly switch between stable epithelial states and motile mesenchymal states to form complex tissue structures.^4^ During these switching events, which are the so-called epithelial to mesenchymal transition (EMT) along with the reverse process called mesenchymal to epithelial transition (MET), the cytoskeletal components are very differently expressed. These different compositions lead to different mechanical and motile behaviors. Besides actin-related structures, crucial elements are keratin IFs, which, with their 37 cytoplasmic members, constitute the largest group of the IF protein-family.^5, 6^ They are typically expressed in epithelial cells in various combinations and provide crucial cell type-specific structural support upon mechanical stresses. These diverse mechanical properties are realized by expressing various specific keratin pairs as keratins obligate heterodimeric complexes representing parallel coiled coils made from two alpha-helical molecules of two sequence-related classes each.^7^ They are also involved in other non-mechanical functions, including regulation of cell growth, migration, and protection from apoptosis.^8, 9^ Several studies have highlighted the potential inhibitory role of keratins for cell mobility and thus the need to downregulate them for EMT to increase cellular mobility. For example, it was found that keratinocytes with all keratins deleted by gene-targeting are softer, more deformable, and more invasive compared to the small overall effect generated after actin depolymerization.^10^ They are also able to migrate twice as fast as wild type cells.^11^ Conversely, keratin re-expression in these studies has the opposite effect. This indicates a direct link between downregulation of keratins during EMT and loss of stiffness with increased migration and invasion ability of tumor cells. Although F-actin networks dramatically reorganize as a prerequisite for morphological and migratory changes during EMT and MET, it is indeed remarkable that the entire IF system is rebuilt by a switch between keratin and vimentin expressions.^12^ As demonstrated for numerous physiological situations, the cytoskeletal systems act synergistically.^13^ By coordinating their functions, they affect each other’s mechanics. However, it is challenging to investigate this mechanical interplay in cells due to their inherent complexity, which makes it difficult to disentangle mechanical crosstalk and regulatory biochemical interactions. In this respect, reconstituted composite cytoskeletal networks have been investigated *in vitro* by quantitative rheological measurements and theoretical modelling. The results have often revealed unexpected mechanical responses. For instance, the presence of microtubules was found to induce unexpected local compressibility^14^ and non-linear strain stiffening in actin networks.^15–17^ Concomitantly, actin reinforces microtubules against compressive loads.^18^

When designing and interpreting such *in vitro* studies, it is important to choose the system parameters carefully as observed when measuring the mechanical properties of composite networks of actin and vimentin. Holding the molecular content constant for different mixing ratios of actin and vimentin has yielded contradictory results for their mechanical response, namely a synergistic increase in the linear elasticity^19^ and formation of stiffer or softer networks depending on the investigation technique.^20, 21^ In this case, however, vimentin filaments carry roughly twice as many monomers per unit length than actin filaments, i.e. total filament length is half compared to that of F-actin. Consequently, it has been shown that actin-vimentin composites with the same total polymer length and mesh size, can be described by a superposition of the mechanical properties of the underlying constituents.^22^

Due to their intrinsic ability to self-assemble *in vitro*, the co-polymerization of actin and keratin is possible without additional accessory proteins. Hence, it has been recently demonstrated that encapsulation of actin and keratin within vesicles leads to the formation of composite networks, where F-actin acts as steric resistance for keratin IFs preventing their collapse to bundled clusters.^23^

Here, we investigate the mechanical properties of *in vitro* reconstituted F-actin and keratin 8/18 (K8-K18) composite networks by bulk shear rheology measurements. We aimed to characterize networks with similar total filament lengths. By systematically varying the relative mass ratios of actin and keratin, we investigate how their relative presence impacts the overall mechanical response of the composites. We complement these rheology measurements with confocal microscopy of the composites.

## Results and discussion

### Co-polymerization and filament length

In order to investigate the mechanical properties of composite networks of actin and K8-K18, we chose the initial concentrations of actin monomers and keratin tetramers that result in comparable total filament lengths of actin and keratin as depicted in Figure 1.

**Fig. 1.**
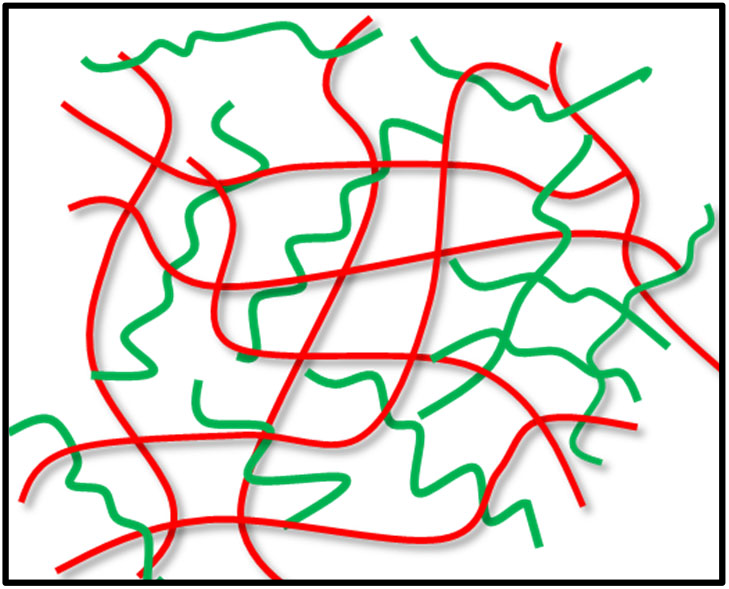
Illustration of an F-actin-keratin composite network with similar total filament length. By co-polymerizing actin monomers at 0.5 mg/ml and keratin tetramers at 0.6 mg/ml, an intermixed network of actin (red) and keratin (green) is formed. At the noted concentrations, networks with the total polymer lengths of the composite will be approximately the same.

We reconstituted composite networks of actin and keratin by co-polymerization of actin monomers and tetrameric complexes of K8 and K18. It is well known that the ionic requirements for polymerization of actin and keratin are quite different. Keratin can assemble *in vitro* into networks in an unusually low ionic strength buffer ^24^ and addition of ions such as Magnesium (Mg^+2^) or potassium (K^+^) at physiological concentrations is enough to induce immediate bundling resulting in heterogeneous networks.^25–27^ In contrast, for actin the addition of mono- and divalent ions to physiological concentrations is essential to initiate the polymerization from globular actin (G-actin) to filamentous actin (F-actin) to form entangled networks.^28^ A striking difference between actin and keratins is the filament persistence length, which is the average length over which filaments stay straight. Keratins have a significantly smaller persistence length (l_p_) than actin, with l_p_ ≈ 0.5 μm and 10 μm, respectively.^29^ Here, we assessed the polymerization efficiency of keratin and actin in different concentrations of MgCl_2_ (1, 0.5, 0.1 and 0 mM) by high-speed sedimentation of protein assemblies followed by SDS-PAGE of pellet and supernatant fractions (data not shown). We found that a lack of Mg^+2^ ions had no effect on the polymerization efficiency of keratin and soluble keratin tetramers were polymerized completely into filaments, whereas for actin at least 1 mM MgCl_2_ was required to ensure complete polymerization. Thus, we had to adjust the polymerization buffer (see Materials and Methods) to a concentration of 1 mM MgCl_2_ for the co-polymerization in order to form actin-keratin networks.

### Linear viscoelasticity of pure actin and keratin networks

Using bulk shear rheology, the linear viscoelastic properties can be quantified by the frequency (ω) dependent complex shear modulus G*(ω) = G′(ω) + iG″(ω), where G′ and G″ are the elastic and viscous moduli, respectively. Under the selected assembly conditions, pure K8-K18 networks at 0.6 mg/ml exhibit predominantly elastic behavior with G′ being consistently larger than G″ by nearly one order of magnitude (Fig. 2*A*). Accordingly, the loss factor tan (ɸ) = G″/G′ has a small value (tan (ϕ) at 1 Hz = 0.13), indicating highly elastic networks. Over the entire frequency range, G′ and G″ show a very weak frequency dependence. A weak power law was obtained by a linear fit of G′ in the log-log-plot yielding a power-law exponent of α(G′) = 0.07. We observed no G″/G′ crossover in the measurable frequency range for pure keratin. This is consistent with results previously reported for K8-K18 network bulk rheological behavior.^24, 30, 31^ By comparison, G′ and G″ of pure F-actin networks showed a more pronounced frequency dependence with α(G′) = 0.23, (Fig. 2 E). Accordingly, the crossover between G′ and G″ regimes was observed at low frequency. Actin networks have a larger loss factor (tan (ϕ) at 1 Hz = 0.52) and, consequently, the elastic contributions are less dominant than in pure keratin networks. These mechanical features are similar to those obtained for entangled actin networks under comparable conditions.^22, 32^

**Fig. 2.**
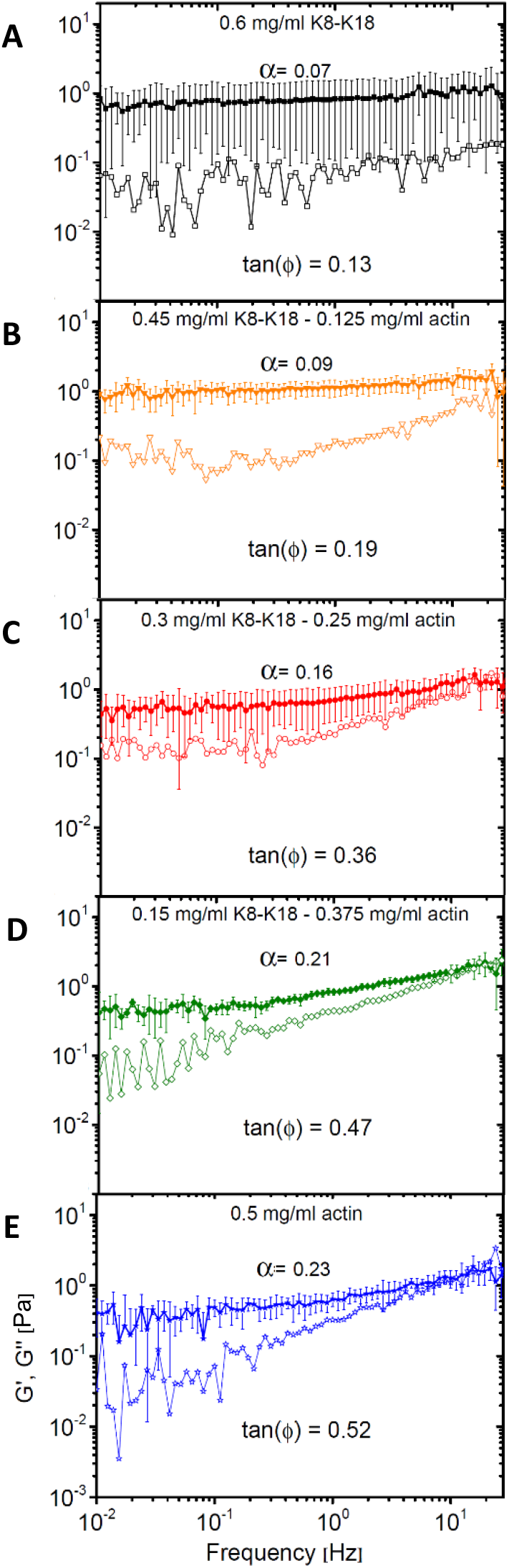
Linear viscoelasticity of reconstituted networks. By comparison, the keratin network (A) shows more predominant elasticity with a weaker frequency dependence of both elastic G′ (filled symbols) and viscous G″ (open symbols) moduli than the F-actin network (E); the keratin network is more elastic (tan ϕ _ker_ = 0.13) than the F-actin network (tan ϕ _act_= 0.52) with no observed G′/G″ crossover, which is a signature of actin networks in this frequency range. Actin-keratin composite networks with varying mixing ratios of actin and keratin (B-D) show intermediate viscoelastic properties. With increasing actin/decreasing keratin content, the frequency dependence of G′ and G″ moduli increases gradually, and the networks become less elastic as the loss factor values increase. Consequently, the G′/G″ crossover appears at high frequency in keratin-dominated networks and shifts gradually to lower frequencies with increasing actin content. α represents the slope between 0.1 and 10 Hz of G′. Data points in all curves represent the mean of at least five independent measurements and error bars represent the standard deviation from the mean.

### Composite networks exhibit an intermediate linear viscoelastic behavior

To date, previous *in vitro* studies have investigated the mechanical properties of only one-component systems of either actin as entangled^33–35^ and crosslinked^36–40^ networks or keratin networks^24, 25, 30, 31, 41^ and single filaments.^42^ In our study, , composite networks of F-actin and K8-K18 with varying mixing ratios revealed an intermediate linear viscoelastic behavior with regard to their composite-specific protein content. With increasing actin/decreasing keratin contents as shown in figure 2 (B-D), the dependence of G′ and G″ on the frequency increased gradually as indicated by the gradual increase in α values; α = 0.09 in keratin-dominated network (0.45 mg/ml K8-K18 – 0.125 mg/ml actin), 0.16 in equal ratio-networks (0.3 mg/ml K8-K18 – 0.25 mg/ml actin) and 0.21 in F-actin-dominated networks (0.15 mg/ml K8-K18 – 0.375 mg/ml actin). Furthermore, the crossover to the predominant viscous regime, observed only in F-actin networks, started to appear in keratin-dominated networks at high frequencies. With increasing actin content, actin contributes more to the composite’s behavior, shifting the crossover point gradually to lower frequencies. The loss factor tan (ϕ) increased with increasing actin/decreasing keratin content, which indicates a smooth transition to less elastic networks, as shown in figure 3A. In figure 3B. We used a Mann-Whitney U test to investigate the statistical significance of the loss factor values with respect to each other. We found p-values less than 0.05, indicating significant difference, for all compared distributions except for the pure networks compared to composites with a respective division of 0.75/0.25 (Table S1, ESI). In figure 3B, we show that the elastic moduli G′ for all networks exhibit only minor variations.

**Fig. 3.**
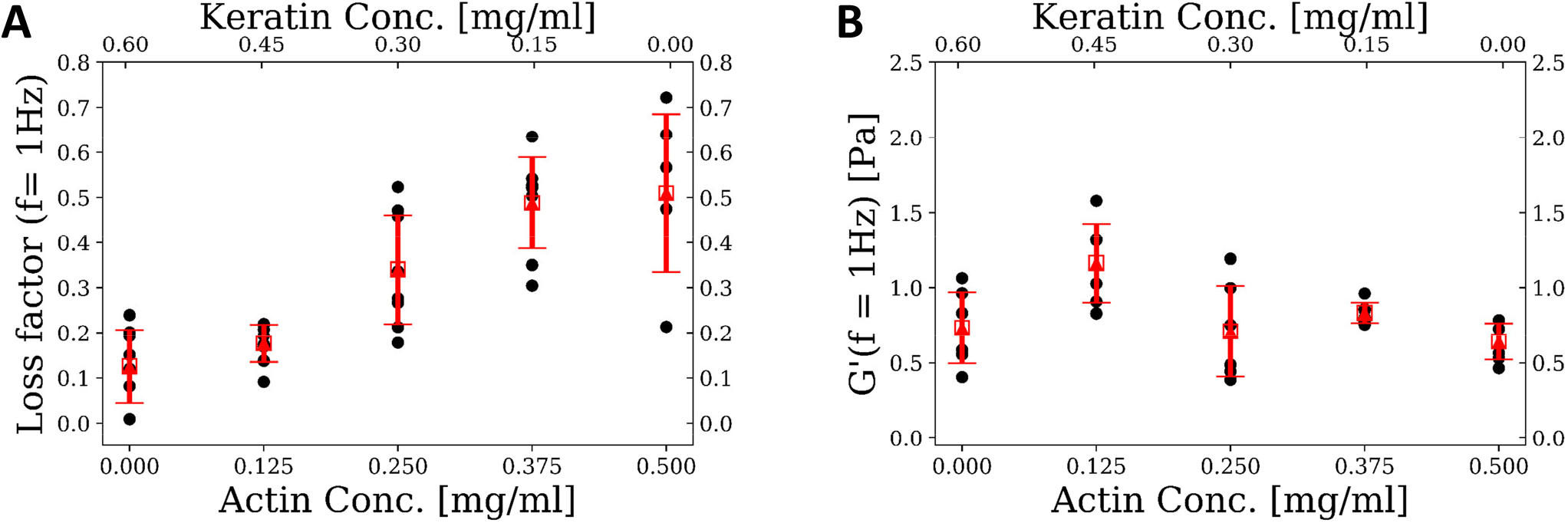
Mechanical properties of pure and composite networks in the linear regime. (A) Mean values of the loss factor tan(ϕ) at f =1Hz of composite networks showing a gradual increase in the loss factor values, i.e. the network’s viscosity, with increasing actin/decreasing keratin content. We found the distributions to be significantly different except for the constellations of pure Actin compared to 0.15 mg/ml Keratin −0.375 mg/ml Actin and pure Keratin compared to 0.45 mg/ml Keratin −0.125 mg/ml Actin respectively. (B) The plateau modulus G0 = G′ (f = 1Hz) for all networks shows only minor variations between networks in the linear regime. In (A) and (B) dots represent single measurements with significance bars in A and error bars (standard deviation from the mean) in B.

Interestingly, important behavioral changes in response to strain were observed only in the non-linear regime as the relative amounts of actin and keratin changed. This situation is more relevant for the physiological conditions as, in general, the cell is exposed to different kinds of stresses and strains and thereby adapts its mechanical properties accordingly.

### Filament-filament interactions constitute a fundamental cause for mechanical responses

The weak power law behavior exhibited in actin networks can be described by the glassy worm-like chain model (GWLC), where the exponent of this power law depends on the level of pre-stress and the interaction strength (ε).^43^ This interaction strength is a phenomenological parameter that is typically neglected in models of entangled networks such as the tube model.^44^ The fundamental idea of the GWLC is an exponential stretching of a wormlike chain’s mode relaxation times (*τ*_*n*_) according to

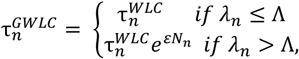

where *λ*_*n*_ = L/n is the half wavelength of eigenmodes with mode number n and *N*_*n*_ = *λ*_*n*_/Λ − 1 the number of interactions per length *λ*_*n*_. Mode relaxation times with *λ*_*n*_ > Λ, where Λ describes a characteristic interaction length, are stretched.^43^ For the interested reader, we would like to refer to the electronic supplementary information (ESI) for a more detailed description of this model. Recently, we demonstrated that this ε can be interpreted as a polymer-specific stickiness and we showed that isotropic networks of K8-K18 assembled in low-salt buffer are much more sticky than F-actin networks assembled in F-buffer.^32^ We used the GWLC model to investigate the non-specific filament-filament interactions that are compiled in this parameter by fitting the expected values obtained from the model to our experimental data (Fig. 4). Using persistence lengths and contour lengths averaged according to the polymer composition, we obtained good fits for every composite, but not for pure keratin networks.

**Fig. 4.**
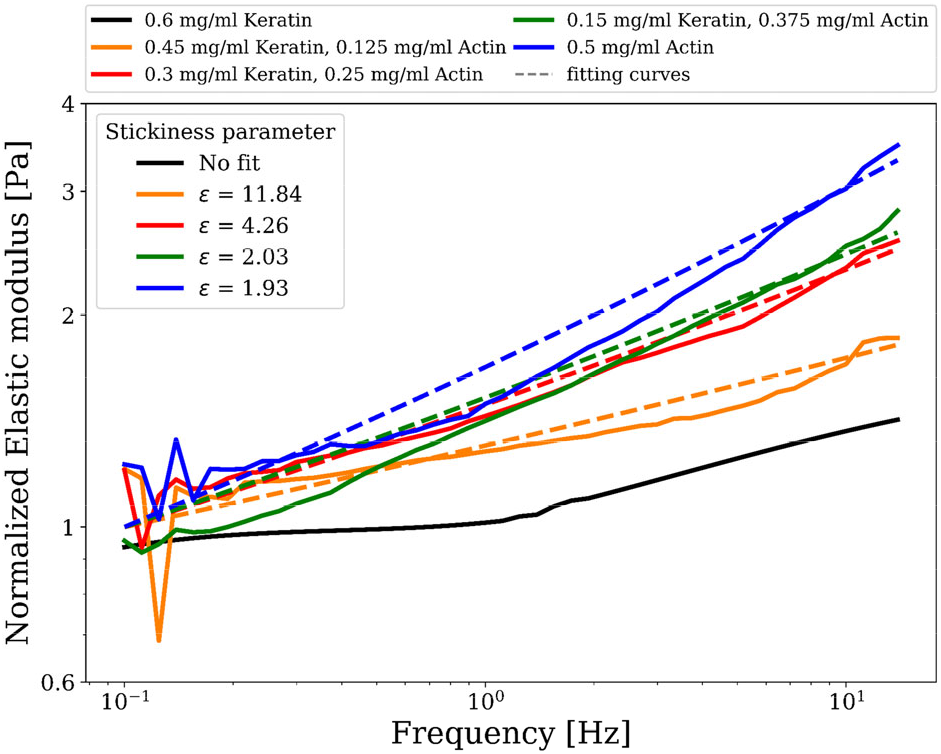
Attractive filament-filament interactions (captured in the stickiness parameter ε) determine the mechanical response of composite networks in linear rheology. Normalized storage moduli (G′) versus frequency for different actin/keratin compositions evaluated with the glassy wormlike chain model. Solid lines represent the measured data and dashed lines represent the fitting curves, yielding values for stickiness ε as shown. For pure keratin, the model could not be fitted to the data.

Fitting the GWLC model to the experimental data yields values for ε that indicate stronger attractive filament-filament interactions for increasing keratin contents in composite networks (Fig. 4). While actin networks are considered a model system for entangled networks, keratin networks form sticky clusters due to pronounced hydrophobic interactions^31, 43^ and, therefore, pure keratin networks are not readily accessible to the GWLC model.

As the GWLC model is based on the assumption of isotropic networks, it cannot capture pure keratin systems due to the inherent bundling under the investigated conditions (Fig. 4, ESI†). Cluster-forming networks may be better described by models for cross-linked networks such as an affine model.^45^ Recently, a detailed model for bundling of keratin was suggested based on the interplay between inter-filament electrostatic and hydrophobic interactions. This model predicts that the process of keratin bundling is determined by the electric charge of the filaments, the number of hydrophobic residues, and the exclusion of the ions from the bundle interior.^46^ Compared with actin, the attractive interactions between filaments in pure keratin structures are much stronger than those in entangled actin networks.^32^ Consequently, the viscous loss modulus in keratin networks during oscillatory shear is significantly reduced compared to the viscous dissipation in actin networks.^47^ Therefore, the viscous modulus G″ of actin networks was observed steeply increasing towards the elastic modulus G′, and the crossover point was observed in the linear regime.

This denotes the transition from a regime dominated by filament interactions within the network to a regime dominated by single filament behavior, which is a signature of actin networks.^48^ The resulting parameters for the stickiness parameter ε are shown in Fig. 4. For increasing keratin content, we observed a stronger attractive interaction between individual filaments, reflected in higher ε values.

### Keratin induces strain stiffening in the non-linear regime

Strain stiffening of biopolymer networks is a feature shared by various intermediate filament proteins.^49^ This physical response is of particular physiological significance for keratin, which makes up a major portion of structural protein found in epithelial cells, and provides protection against large-scale deformation. Interestingly, keratin networks exhibit strain stiffening even in the absence of cross-linkers or divalent cations,^24, 31^ while in actin networks the strain stiffening depends on the cross-links^48, 50^.

These features of keratin and actin assemblies are reflected in the differential shear modulus K for the different composite networks. For networks with lower keratin content, we see weak strain-stiffening effects comparable to that of pure actin networks. This is expressed in a maximum value K that is two orders of magnitude lower than for pure keratin and keratin-dominated networks, (Fig. 5, inset) appearing at stresses shifted to values more than one order of magnitude lower (Fig. 5).

**Fig. 5.**
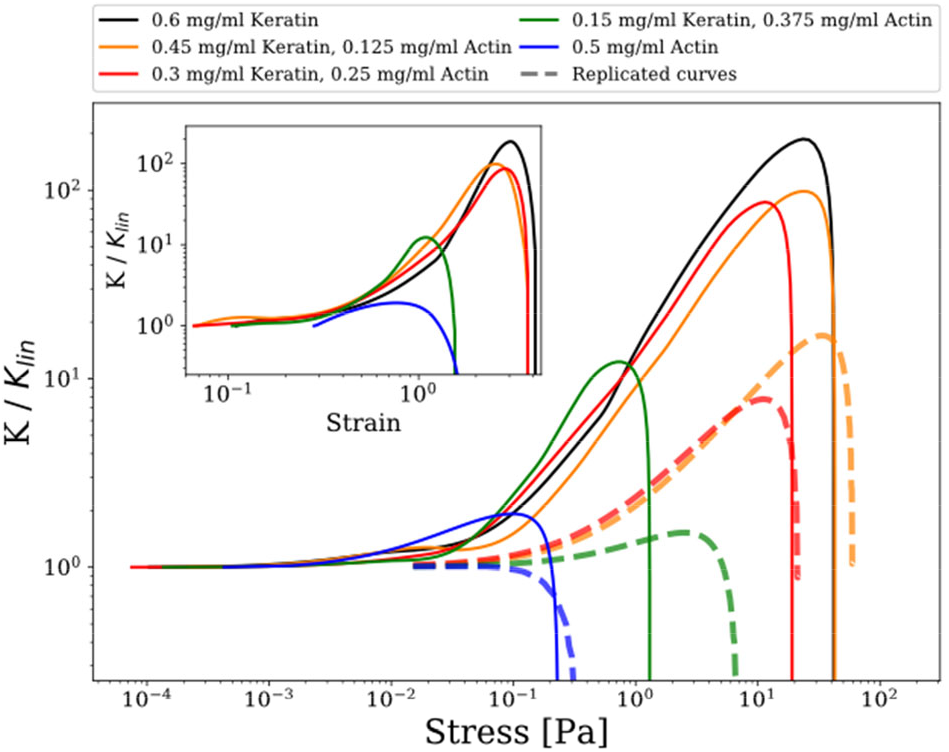
Actin-keratin composite networks exhibit strain stiffening increasing in manifestation with increasing keratin content. This is expressed in the differential shear modulus K = *dσ/dγ* rescaled by its value in the linear regime K_*lin*_ and plotted versus stress *σ* for comparison to the prediction of the model. Solid lines are measurement curves and dotted lines are replicated curves with the non-linear extension of the GWLC. The replicated curves are an order of magnitude lower than the measured data, but show a comparable trend. The inset shows K versus strain

In the frame of the EMT, keratins are down-regulated, and mesenchymal cells do not strain stiffen and can readily migrate. In contrast, high keratin content improves strain stiffening which helps non-motile epithelial cells to remain stable and to keep tissue formations intact.

The experimental curves can be replicated qualitatively in the frame of the GWLC using the parameters obtained from fitting the model to the linear data G*(f) (Fig. 5). Even though the measured maximum differential modulus is one order of magnitude higher and shifted to lower stresses in the composite networks, the model curves represent the same qualitative features and the same overall trend to express an increasing strain-stiffening at larger strains/stresses for higher keratin contents. This underestimation of K_*max*_ has been described previously by Golde *et al.*^32^ as a limiting case for the applicability of the GWLC giving rise to the assumption that keratin networks, due to strong filament interactions in combination with a small persistence length, express a phenomenological behavior that is more consistently described as crosslinked rather than entangled.^51^

### Network architectures

Using spinning disc confocal microscopy, we examined the architecture of all networks (10% labeled sample) under these assembly conditions. As mentioned above, keratin is known to quickly form networks due to its high self-affinity, and bundled networks if positively charged ions are present.^24, 27^ We initiated the assembly at low temperature by mixing the protein solution with the assembly buffer on ice in order to slow down the assembly process. A dense, heterogeneous network was obtained as shown in figure 6A. Interestingly, a bundled keratin network was also formed even at a very low protein concentration (e.g. as low as 0.1 mg/ml) (Fig. S1A, ESI†). In the absence of salts, keratin can assemble into isotropic networks even at high protein concentrations as shown in Fig. S1B ESI†. By contrast, actin at 0.5 mg/ml assembled solely into highly isotropic entangled networks under selected ionic conditions (Fig. 6B). Actin filaments often associate into bundles and networks with diverse structures only in the presence of ‘’bundling factors’’ such as high salt conditions or actin-binding proteins.^52–54^ Consequently, bundling of actin was not an issue, and actin filaments completely arranged into isotropic networks under the investigated conditions. In composite networks, actin provides steric hindrance, creating “obstacles” in between keratin filaments, thereby preventing or reducing their tight bundling. This observation is consistent with a previous *in vitro* study on actin-keratin composites encapsulated in vesicles.^23^ However, we observed that this steric effect became less pronounced in increasingly keratin-dominated networks (Fig. 6C). Under these conditions, actin and keratin were observed completely co-localized, but mostly appeared as a large cluster. In actin-dominated regimes, as well as equal ratio networks, keratin appeared as extended networks as well as very small clusters surrounded by homogenous actin networks (Fig. 6D and E). In all composites, both keratin and actin networks did not demix, but were rather observed as a composite system with networks of interdependent architectures.

**Fig. 6.**
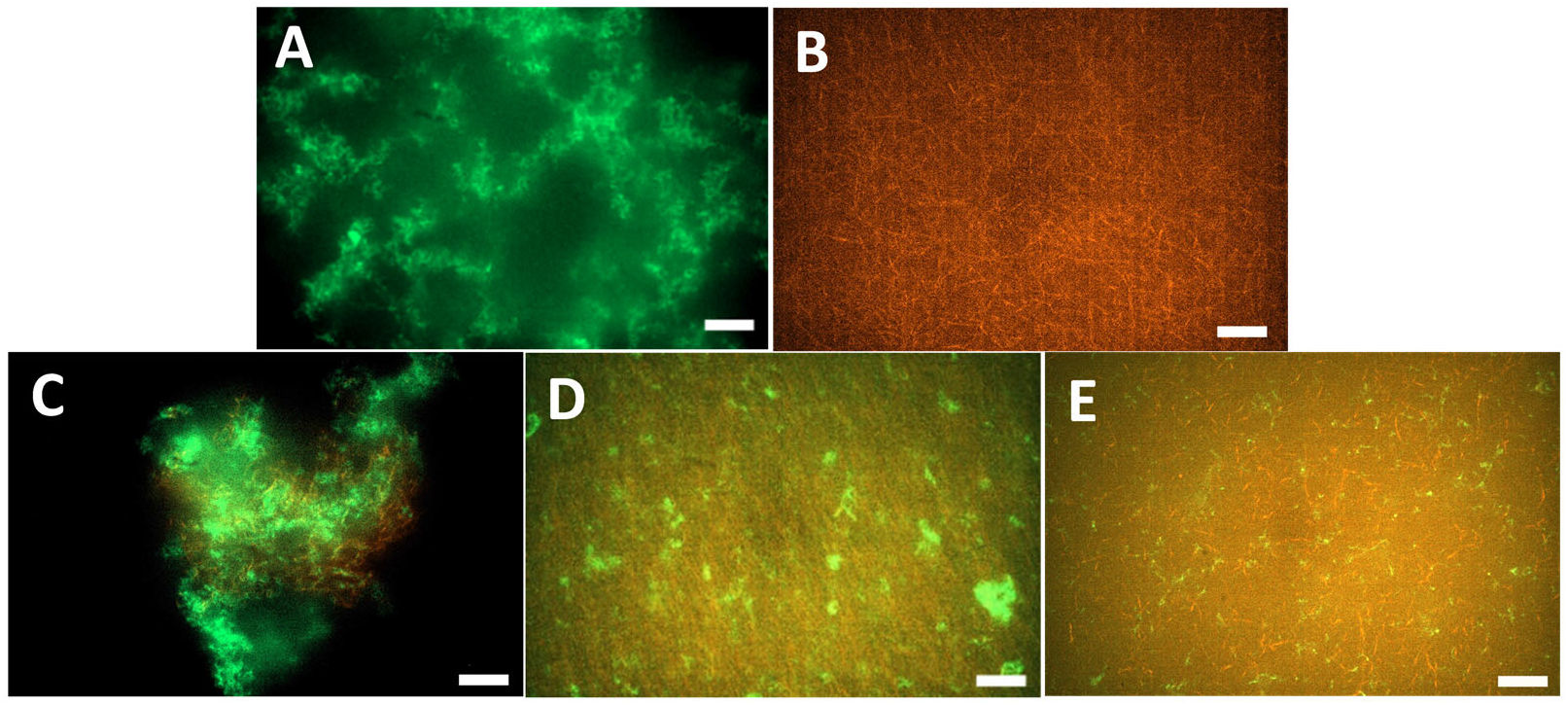
Confocal micrographs of in situ formed networks. Keratin assembles into a dense heterogeneous network immediately after the addition of F-buffer (A), while actin assembles solely into an isotropic entangled network (B). In keratin-dominated networks, a complete co-localization of F-actin and keratin networks culminates in a large cluster (C). In equal ratio and F-actin-dominated networks, the isotropic actin networks provide steric hindrance against tight bundling of keratin, and thus keratin appears as filaments and small clusters in these composites (D and E respectively). Scale bar for all images: 10 μm.

To visualize keratin networks, we developed a new method for labeling the wild type keratin without mutations (described in Materials and Methods). As a control, we tested the bulk mechanical properties of the 10% labeled keratin samples. We found that these labeled samples behave as unlabeled keratin, indicating that the filaments and network properties were not affected by the presence of 10% labeled keratin (Fig.S2, ESI†).

## Experimental

### Polymer lengths

The mass per unit length (m_L_) for actin is 2.66×10^−11^ g/m^55^, and for K8-K18 in Tris-based assembly buffer is 3.16×10^−11^ g/m^24^. At 0.5 mg/ml actin monomers (11.9 μM) and 0.6 mg/ml K8-18 tetramers (11.34 μM), both actin and keratin networks will have similar total filament length. Using these concentrations as our boundary conditions, we can select actin-keratin mixtures with the same total polymer length per unit volume.

### Protein preparation and co-polymerization

G-actin was prepared from rabbit muscle and stored at −80 °C in G-Buffer (2 mM Tris-HCl pH 7.5, 0.2 mM ATP, 0.1 mM CaCl_2_, 1 mM DTT, 0.01% NaN_3_, pH 7.8) as described previously.^56^ Vectors containing keratin genes (pET 24a-K8 and pET 23a-K18) were transformed into **E**. coli BL21 for protein expression. Recombinant K8 and K18 were isolated and purified as previously described.^57^

For reconstitution, purified K8 and K18 proteins were mixed in equimolar amounts and renatured by stepwise dialysis against 8 M urea, 2 mM Tris–HCl, 1 mM DTT, pH 9.0 with stepwise reduction of urea concentration (6 M, 4 M, 2 M, 1 M and 0 M). Each dialysis step was done for 20 minutes at room temperature, then the dialysis was continued overnight against 2 mM Tris–HCl, 1 mM DTT, pH 9.0 at 4 °C. The final protein concentration was determined by measuring the absorption at 280 nm using a DU 530 UV/vis Spectrophotometer (Beckman Coulter Inc., USA). Composite networks were prepared by mixing actin monomers at 0.5 mg/ml and K8-K18 tetramers at 0.6 mg/ml in varying mixing ratios. Assembly of pure and mixed networks was initiated by adding 1/10 volume of 10x F-buffer (20 mM Tris-HCl, 1 M KCl, 10 mM MgCl_2_, 2 mM ATP, 10 mM DTT, pH 7.5) to the protein sample.

### Shear rheology

Rheology measurements were conducted on a strain-controlled ARES rheometer (TA Instruments, USA) using 40 mm plate-plate geometry with 140 μm gap width at 20 °C. In all measurements, F-buffer was mixed with the protein mixture on ice, and a sample volume of 200 μl was loaded quickly between the two plates. F-Buffer with the same conditions as in the sample was distributed around the sample to prevent artifacts from interfacial elasticity and a solvent trap was placed around the sample to prevent evaporation. The sample was given two hours to equilibrate between the rheometer plates as the filament assembly was monitored with a dynamic time sweep for 2 hours, one data point per minute at a frequency (ω) of 1 Hz and a strain (γ) of 2%. The linear viscoelastic response of equilibrated networks was measured with dynamic frequency sweeps ranging from 0.01 Hz to 80 Hz at a strain of 2% and 20 points per decade. Data points plotted in figure 2 are the mean of at least five independent rheology measurements. A transient step rate test with a strain rate of 0.1 s^−1^ was applied to measure the strain-dependent stress in the non-linear strain regime. The differential shear modulus (K) was determined with a self-written Python script, calculating the gradient of the smoothed stress data divided by the strain step width. The linear differential shear modulus K_*lin*_ is given by the first non-negative value of the smoothed stress data, whereas negative stress is measured due to technical limitations, particularly for small strains, lacking every physical relevance. The linear frequency sweeps as well as the nonlinear step rate measurements were evaluated in the frame of the GWLC model with a self-written Python script. At least six independent rheology measurements were taken into account for the evaluation and comparison with the model predictions. Details about the model are presented in the Supplemental Material.

### Protein labelling, sample preparation and imaging

We have developed a new method for direct labelling of the wild type K8-K18 without mutation, unlike the techniques used in previous studies.^23, 42^ The labelling is based on the coupling of Atto-488-NHS-ester (ATTO-TEC, Siegen, Germany) to the free amine (NH_2_) residues (Lysine or terminus) in K8 tetramers. K8 was renatured by stepwise dialysis against the labelling buffer (2 mM sodium phosphate, pH 8.5) as described above. K8 tetramers were mixed with Atto-488-NHS-ester solubilized in DMSO at 10 mM at a molar ratio of 5:1. The mixture was incubated in the dark for 1 hour at room temperature. In one step, the free dye was removed, and labelled K8 was denatured into monomers by overnight dialysis against 8M urea, 2 mM sodium phosphate buffer pH 8.5 at 4 °C. On the second day, the labelling buffer was exchanged by the dialysis buffer (8 M urea, 2 mM Tris-HCl, pH 9.0) by dialysis for 2 hours at room temperature. The concentration of labelled K8 was determined by measuring the absorbance at 280 and 500 nm using a DU 530 UV/vis Spectrophotometer (Beckman Coulter Inc., USA). Labelled K8 was then distributed to appropriate aliquots and stored at −80 °C. One day before imaging, equal amounts of 20% labelled K8 were mixed with unlabelled K8 and K18 in 8 M urea, 2 mM Tris-HCl, pH 9.0 to have a final sample labelling of 10% by concentration. This mixture was renatured into tetramers by stepwise dialysis against 2 mM Tris-HCl, pH 9.0.

For network visualization, labelled K8-K18 prepared in the previous step was mixed with unlabelled K8-K18 tetramers in a ratio of 1:10, with a final protein concentration of 0.6 mg/ml. Fluorescently labelled actin was prepared by mixing G-actin at 5 μM with phalloidin–tetramethylrhodamine B isothio-cyanate (phalloidin– TRITC – Sigma-Aldrich Co.) in a molar ratio of 1:1. For composite networks, 10% labelled keratin samples were mixed with labelled actin samples in varying ratios. Networks were created by adding 1/10 volume fraction of 10x F-buffer to the sample and immediately pipetting it into an experimental sample chamber to rest for 2 hours at room temperature before imaging as described in ^36^. Images were captured using a spinning disc confocal microscope (inverted Axio Observer.Z1/Yokogawa CSU-X1A 5000, Carl Zeiss Microscopy GmbH, Germany), 100x oil immersion objective NA 1.40 with a Hamamatsu camera at an exposure time of 100 ms.

## Discussion and Conclusions

To study how actin and keratins affect each other’s organization, we have developed a suitable labelling procedure for K8-K18, which enabled us to find that actin sterically hinders and effectively reduces the tight bundling of keratin in some composites. This observation highlights the supportive role of actin within the cell as it enables keratin networks to extend and provide protection to the entire cell. Previous reports demonstrate that the downregulation of keratins can provide space in the cell periphery to enable F-actin networks to reorganize more freely to form protrusions for migration. The presence of an intact keratin network in the cell periphery, however, slows down actin reorganization.^58, 59^ These results support our observation that this hindrance effect provided by actin diminishes in keratin-dominated networks.

We investigated the bulk rheology of F-actin/K8-K18 filament composite networks with varying relative concentrations. This may provide insights into the mechanical and structural interplay they might display in a physiological situation, for instance, for cells integrated into an epithelial cell layer when starting to turn into a migrating cell during and after EMT. In the latter situation, cells build up a vimentin IF system, which provides completely different mechanical properties.^60–62^ Hence, this change of the IF system is probably of direct importance to the process as vimentin IF provides a much softer counterpart system to F-actin with strain stiffening more than one order of magnitude lower than K8-K18 IF.^22^

Here, we show that for small deformations, i.e., the linear deformation regime, the composites revealed an intermediate linear viscoelastic behavior compared to that of the pure networks. This provides important new information about the mechanical coupling between networks of these two structural proteins within the cell environment. Similar to mixed F-actin/vimentin IF networks, the mechanics of composite F-actin/K8-K18 filament networks can be described from their respective substructures as a superposition in the frame of the GWLC model.^22^ This suggests that cells can tune their network mechanics by varying the relative ratios of actin and keratin. They may likewise utilize this tunability when changing their viscoelastic properties by varying cytoskeletal network components to meet the mechanical demands in various different simple epithelia such as found in lung and small intestine. These changes are especially important for large deformations, i.e. non-linear deformations. The F-actin/K8-K18 filament composites show drastic strain stiffening under larger deformations, which is induced and dominated by the keratin content. Strain stiffening can be especially used to absorb large external forces, ensuring that tissue structures such as the epithelium remain stable. This non-linear behavior is relevant for cells migrating through dense tissues as they are subjected to different mechanical loads. This may explain the physiological necessity to downregulate keratins which is accompanied by cell softening and enhanced migration ability. Our rheological measurements suggest that the downregulation of K8-K18 may contribute to the loss of cell stiffness as needed for migration. This is supported by several knockout experiments showing that cells lacking keratins are more deformable and invasive, ^10^ migrating faster than wild type cells.^11^ These effects are also much more pronounced than the softening effects resulting from actin depolymerization.^10^ In particular, loss of K8-K18 in epithelial cancer cells was found to increase collective cell migration^63^. Interestingly, the mesenchymal IF vimentin was also often found co-localized with keratin in highly motile epithelial cells.^64, 65^ Vimentin is thus proposed to have a function in migrating cells which may be relevant for that of keratin. In a previous study, it was suggested that vimentin templates short-lived microtubule formation, as vimentin has a comparatively long half-life.^66^ Vimentin templating seems to persist in order to support the directed extension of microtubules, thereby improving persistence of cell migration and enhancing migration efficiency. This behaviour, together with the loss of keratins, elevates the speed of cell migration while reducing persistence. By contrast, increased keratin expression has the opposite impact on cell migration.

The behavior of composite cytoskeletal networks in cells is also highly affected by different cross-linkers. For example, many actin cross-linkers mediate actin bundles in cells^67^ and induce strain stiffening in *in vitro* reconstituted actin networks.^48, 50^ In the case of keratin, depletion of keratin-associated plectin, which cross-bridges individual IFs and connects them to other cytoskeletal components, alters the organization and dynamics of keratin, but does not affect the overall mechanical properties of the cell.^68, 69^ Further studies using biological cytolinkers that crosslink keratin to actin are necessary to help in understanding the complex behavior of this composite.

In summary, our findings suggest that the up- and downregulation of keratin is a physiological tool of cells to modulate their strain stiffening behavior for different physiological situations. The basic mechanical requirements for a cell fixed within an epithelium versus those migrating within tissues, as extensively encountered during embryogenesis, is laid down in the developmental building plan of the respective tissue. Forming complex tissues highly relies on stable stationary cell states (epithelial cells) as well as migratory cells (mesenchymal) for specialization, which themselves are accompanied with respective up- or downregulation of keratin and vimentin. Further fine-tuning may then be achieved by recruiting associated proteins and by posttranslational modifications.^3, 70, 71^

## Acknowledgements

We acknowledge funding by the ESF: European Social Fund for I.E. (ESF—100327895), P.M. (ESF—100316844), and C.T. (ESF—100380880). Furthermore, we acknowledge funding by the European Research Council (ERC-741350) and the German Research Foundation (INST 268/296-1 FUGG & HE 1853/11-1).

## References

1. D. A. Fletcher and R. D. Mullins, Nature, 2010, 463, 485–492.

2. T. Hohmann and F. Dehghani, Cells, 2019, 8, 362.

3. F. Huber, J. Schnauss, S. Ronicke, P. Rauch, K. Muller, C. Futterer and J. Kas, Adv. Phys., 2013, 62, 1–112.

4. D. H. Kim, T. Xing, Z. Yang, R. Dudek, Q. Lu and Y.-H. Chen, J. Clin. Med., 2018, 7, 1.

5. H. Herrmann and U. Aebi, Cold Spring Harb. Perspect. Biol., 2016, 8, a018242.

6. J.-F. Nolting and S. Köster, New J. Phys. , 2013, 15, 045025.

7. J. T. Jacob, P. A. Coulombe, R. Kwan and M. B. Omary, Cold Spring Harb. Perspect. Biol., 2018, 10, a018275.

8. S. Yoon and R. E. Leube, Essays Biochem., 2019, 63, 521–533.

9. X. Pan, R. P. Hobbs and P. A. Coulombe, Curr. Opin. Cell Biol., 2013, 25, 47–56.

10. K. Seltmann, A. W. Fritsch, J. A. Kas and T. M. Magin, PNAS, 2013, 110, 18507–18512.

11. K. Seltmann, W. Roth, C. Kroger, F. Loschke, M. Lederer, S. Huttelmaier and T. M. Magin, J. Invest. Dermatol., 2013, 133, 181–190.

12. B. O. Sun, Y. Fang, Z. Li, Z. Chen and J. Xiang, Biomed. Rep., 2015, 3, 603–610.

13. M. Schoumacher, R. D. Goldman, D. Louvard and D. M. Vignjevic, J. Cell Biol., 2010, 189, 541–556.

14. V. Pelletier, N. Gal, P. Fournier and M. L. Kilfoil, Phys. Rev. Lett, 2009, 102, 188303.

15. Y.-C. Lin, G. H. Koenderink, F. C. MacKintosh and D. A. Weitz, Soft Matter, 2011, 7, 902–906.

16. S. N. Ricketts, J. L. Ross and R. M. Robertson-Anderson, Biophys. J., 2018, 115, 1055–1067.

17. F. Burla, Y. Mulla, B. E. Vos, A. Aufderhorst-Roberts and G. H. Koenderink, Nature, 42254, 019–0036.

18. C. P. Brangwynne, F. C. MacKintosh, S. Kumar, N. A. Geisse, J. Talbot, L. Mahadevan, K. K. Parker, D. E. Ingber and D. A. Weitz, J. Cell Biol., 2006, 173, 733–741.

19. O. Esue, A. A. Carson, Y. Tseng and D. Wirtz, J Biol Chem, 2006, 281, 30393–30399.

20. M. H. Jensen, E. J. Morris, R. D. Goldman and D. A. Weitz, Bioarchitecture, 2014, 4, 138–143.

21. H. Lopez-Menendez and L. Gonzalez-Torres, J. Mech. Phys. Solids, 2019, 127, 208–220.

22. T. Golde, C. Huster, M. Glaser, T. Handler, H. Herrmann, J. A. Kas and J. Schnauss, Soft Matter, 2018, 14, 7970–7978.

23. J. Deek, R. Maan, E. Loiseau and A. R. Bausch, Soft Matter, 2018, 14, 1897–1902.

24. P. Pawelzyk, H. Herrmann and N. Willenbacher, Soft Matter, 2013, 9, 8871.

25. A. Leitner, T. Paust, O. Marti, P. Walther, H. Herrmann and M. Beil, Biophys. J., 2012, 103, 195–201.

26. I. Martin, M. Moch, T. Neckernuss, S. Paschke, H. Herrmann and O. Marti, Soft Matter, 2016, 12, 6964–6974.

27. C. Y. Hemonnot, M. Mauermann, H. Herrmann and S. Koster, Biomacromolecules, 2015, 16, 3313–3321.

28. H. Lodish, A. Berk, S. L. Zipursky, P. Matsudaira, D. Baltimore and J. Darnell, Molecular Cell Biology, 2000.

29. A. F. Pegoraro, P. Janmey and D. A. Weitz, Cold Spring Harb. Perspect. Biol., 2017, 9, a022038.

30. S. Yamada, D. Wirtz and P. A. Coulombe, Mol. Biol. Cell, 2002, 13, 382–391.

31. P. Pawelzyk, N. Mücke, H. Herrmann and N. Willenbacher, PLoS ONE, 2014, 9.

32. T. Golde, M. Glaser, C. Tutmarc, I. Elbalasy, C. Huster, G. Busteros, D. M. Smith, H. Herrmann, J. A. Käs and J. Schnauß, Soft Matter, 2019, 15, 4865–4872.

33. B. J. Gurmessa, N. Bitten, D. T. Nguyen, O. A. Saleh, J. L. Ross, M. Das and R. M. Robertson-Anderson, Soft Matter, 2019, 15, 1335–1344.

34. M. L. Gardel, M. T. Valentine, J. C. Crocker, A. R. Bausch and D. A. Weitz, Phys. Rev. Lett., 2003, 91, 158302.

35. T. T. Falzone and R. M. Robertson-Anderson, ACS Macro Lett., 2015, 4, 1194–1199.

36. D. Strehle, J. Schnauss, C. Heussinger, J. Alvarado, M. Bathe, J. Kas and B. Gentry, Eur. Biophys. J., 2011, 40, 93–101.

37. B. Gurmessa, S. Ricketts and R. M. Robertson-Anderson, Biophys. J., 2017, 113, 1540–1550.

38. M. L. Gardel, J. H. Shin, F. C. MacKintosh, L. Mahadevan, P. Matsudaira and D. A. Weitz, Science, 2004, 304, 1301–1305.

39. D. Strehle, P. Mollenkopf, M. Glaser, T. Golde, C. Schuldt, J. A. Kas and J. Schnauss, Molecules, 2017, 22, 1–11.

40. R. Tharmann, M. M. Claessens and A. R. Bausch, Phys. Rev. Lett., 2007, 98, 088103.

41. S. Yamada, D. Wirtz and P. A. Coulombe, J. Struct. Biol., 2003, 143, 45–55.

42. C. Lorenz, J. Forsting, A. V. Schepers, J. Kraxner, S. Bauch, H. Witt, S. Klumpp and S. Koster, Phys. Rev. Lett., 2019, 123, 188102.

43. K. Kroy and J. Glaser, New. J. Phys., 2007, 9, 416–416.

44. H. Isambert and A. C. Maggs, Macromolecules, 1996, 29, 1036–1040.

45. F. C. MacKintosh, J. Kas and P. A. Janmey, Phys. Rev. Lett., 1995, 75, 4425–4428.

46. E. Haimov, R. Windoffer, R. E. Leube, M. Urbakh and M. M. Kozlov, Biophys. J., 2020, 119, 65–74.

47. O. Lieleg, M. M. Claessens, Y. Luan and A. R. Bausch, Phys. Rev. Lett., 2008, 101, 108101.

48. C. Semmrich, T. Storz, J. Glaser, R. Merkel, A. R. Bausch and K. Kroy, PNAS, 2007, 104, 20199–20203.

49. C. Storm, J. J. Pastore, F. C. MacKintosh, T. C. Lubensky and P. A. Janmey, Nature, 2005, 435, 191–194.

50. M. L. Gardel, J. H. Shin, F. C. MacKintosh, L. Mahadevan, P. Matsudaira and D. A. Weitz, Science, 2004, 304, 1301–1305.

51. C. Broedersz and F. MacKintosh, Soft Matter, 2011, 7, 3186–3191.

52. J. X. Tang and P. A. Janmey, Biol. Bull., 1998, 194, 406–408.

53. C. Schuldt, J. Schnauss, T. Handler, M. Glaser, J. Lorenz, T. Golde, J. A. Kas and D. M. Smith, Phys. Rev. Lett., 2016, 117, 197801.

54. J. Schnauß, T. Golde, C. Schuldt, B. S. Schmidt, M. Glaser, D. Strehle, T. Händler, C. Heussinger and J. A. Käs, Phys. Rev. Lett., 2016, 116, 108102.

55. U. Aebi, R. Millonig, H. Salvo and A. Engel, Ann. N. Y. Acad. Sci., 1986, 483, 100–119.

56. B. Gentry, D. Smith and J. Kas, Phys. Rev. E. , 2009, 79, 031916.

57. K. L. A. U. Herrmann Hh, Methods Cell. Biol., 2004, 78, 3–24.

58. A. W. Holle, M. Kalafat, A. S. Ramos, T. Seufferlein, R. Kemkemer and J. P. Spatz, Sci. Rep., 2017, 7, 45152.

59. S. Karsch, F. Buchau, T. M. Magin and A. Janshoff, Cell. Mol. Life Sci., 2020, 77, 4397–4411.

60. A. E. Patteson, A. Vahabikashi, K. Pogoda, S. A. Adam, K. Mandal, M. Kittisopikul, S. Sivagurunathan, A. Goldman, R. D. Goldman and P. A. Janmey, J. Cell Biol., 2019, 218, 4079–4092.

61. A. Aufderhorst-Roberts and G. H. Koenderink, Soft Matter, 2019, 15, 7127–7136.

62. J. Block, H. Witt, A. Candelli, E. J. Peterman, G. J. Wuite, A. Janshoff and S. Koster, Phys. Rev. Lett., 2017, 118, 048101.

63. A.-M. Fortier, E. Asselin and M. Cadrin, J. Biol. Chem., 2013, 288, 11555–11571.

64. B. M. Chung, J. D. Rotty and P. A. Coulombe, Curr. Opin. Cell Biol., 2013, 25, 600–612.

65. C. Velez-delValle, M. Marsch-Moreno, F. Castro-Munozledo, I. J. Galvan-Mendoza and W. Kuri-Harcuch, Sci. Rep., 2016, 6, 24389.

66. Z. Gan, L. Ding, C. J. Burckhardt, J. Lowery, A. Zaritsky, K. Sitterley, A. Mota, N. Costigliola, C. G. Starker, D. F. Voytas, J. Tytell, R. D. Goldman and G. Danuser, Cell Syst., 2016, 3, 252–263 e258.

67. S. J. Winder and K. R. Ayscough, J. Cell Sci., 2005, 118, 651–654.

68. M. Moch, R. Windoffer, N. Schwarz, R. Pohl, A. Omenzetter, U. Schnakenberg, F. Herb, K. Chaisaowong, D. Merhof, L. Ramms, G. Fabris, B. Hoffmann, R. Merkel and R. E. Leube, PLoS ONE, 2016, 11, e0149106.

69. N. Bonakdar, A. Schilling, M. Sporrer, P. Lennert, A. Mainka, L. Winter, G. Walko, G. Wiche, B. Fabry and W. H. Goldmann, Exp. Cell Res., 2015, 331, 331–337.

70. M. P. Serres, M. Samwer, B. A. Truong Quang, G. Lavoie, U. Perera, D. Gorlich, G. Charras, M. Petronczki, P. P. Roux and E. K. Paluch, Dev. Cell, 2020, 52, 210–222 e217.

71. Y. Mulla, F. C. MacKintosh and G. H. Koenderink, Phys. Rev. Lett., 2019, 122, 218102.

