## Supplementary material for "Keratins determine network stress responsiveness in reconstituted actin-keratin filament systems": ESI..docx

**The glassy wormlike chain model.** The GWLC model used for this study has been comprehensively described previously.^1-3^ The GWLC is an extension of the wormlike chain (WLC) for semiflexible polymer networks. The interactions of a test chain with its environment is taken into account by stretching the mode relaxation spectrum of the WLC exponentially. The mode relaxation times of all eigenmodes of (half-) wavelength $\lambda_{n}$ = L/n and mode number n for a WLC with persistence length $l_{p}$ and the transverse drag coefficient $\zeta_{\perp}$ are given by

$$\tau_{n}^{WLC}= \zeta_{\perp} / (\frac{l_{p} k_{B} T \pi^{4}}{\lambda_{n}^{4}}+f \pi^{2}/\lambda_{n}^{2})$$

The relaxation times of the GWLC are modified according to

$$\tau_{n}^{GWLC}= \left\{ \begin{aligned} \tau_{n}^{WLC} if \lambda_{n}\leq\Lambda\\ \tau_{n}^{WLC}e^{{\varepsilon N}_{n}} if \lambda_{n}>\Lambda, \end{aligned} \right.$$

where $N_{n}$ = $\lambda_{n}$/ Λ − 1 is the number of interactions per length $\lambda_{n}$ with the average distance between interaction points, L the contour length of the test filament, the stretching parameter ε controlling how strong the modes are slowed down by interactions with its environment, and $f$ that describes a homogeneous backbone tension accounting for existing pre-stress. The complex linear shear modulus in the high-frequency regime is given by

*Equation 1*

$$G^{*}\left( \omega\right)= \Lambda/(5\xi^{2}\chi(\omega))$$

where $\xi$ is the mesh size of the network. The micro-rheological, linear response function $\chi(\omega)$ to a point force at the ends of the GWLC is calculated as

$$\chi\left( \omega\right)=\frac{L^{4}}{\pi^{4} l_{p}^{2} k_{B} T} \sum_{n=1}^{\infty} \frac{1}{(n^{4}+n^{2}f/f_{E})(1+i\omega\tau_{n}^{GWLC}/2)}$$

with the Euler buckling force $f_{E}=l_{p}k_{B}T\pi^{2}/L^{2}$. For the linear regime, $f$ is set to zero.

In the nonlinear regime, the differential shear modulus K = dσ/dγ is approximated via (Equation 1) at a constant frequency as a function of the backbone tension$f$:

$$K\left( f \right)=|G_{\omega}^{*}|(f)$$

Where $f$ is related to the macroscopic stress σ via f = 5σ$\xi^{2}$. The effect of pre-stress on the stretching parameter is introduced via a linear barrier height reduction:

$$\varepsilon\to\varepsilon-f\delta/k_{B}T$$

where δ should be interpreted as an effective width of a free energy well. The mean values of ξ, Λ, and ε obtained from fitting the linear regime for each polymer type were used to replicate the measured curves. δ was used as the only free parameter to effectively shift the peak of K both in terms of σ and the maximum value $K_{max}$.

Fig. S1. Confocal micrographs of in situ formed keratin networks assembled in different salt conditions. (A) In F-buffer, a bundled network is formed even at low protein concentration (0.1 mg/ml). (B) In standard low-salt buffer (10 mM Tris PH 7), an isotropic network is formed.

**A**

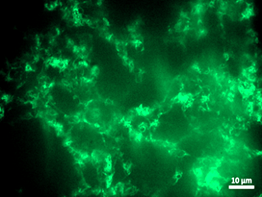

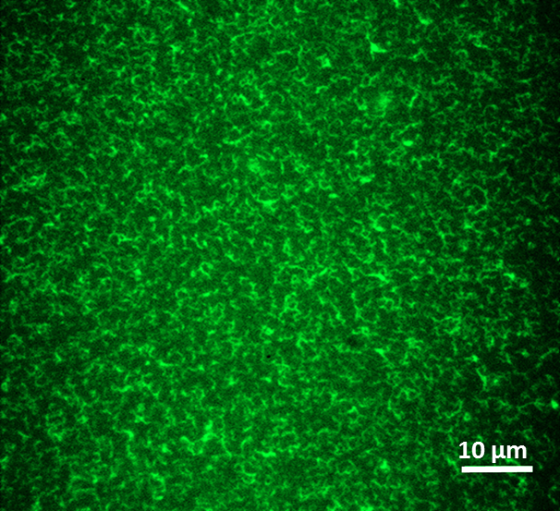

**B**

Fig. S1. Confocal micrographs of in situ formed keratin networks assembled in different salt conditions. (A) In F-buffer, a bundled network is formed even at low protein concentration (0.1 mg/ml). (B) In standard low-salt buffer (10 mM Tris PH 7), an isotropic network is formed.

| 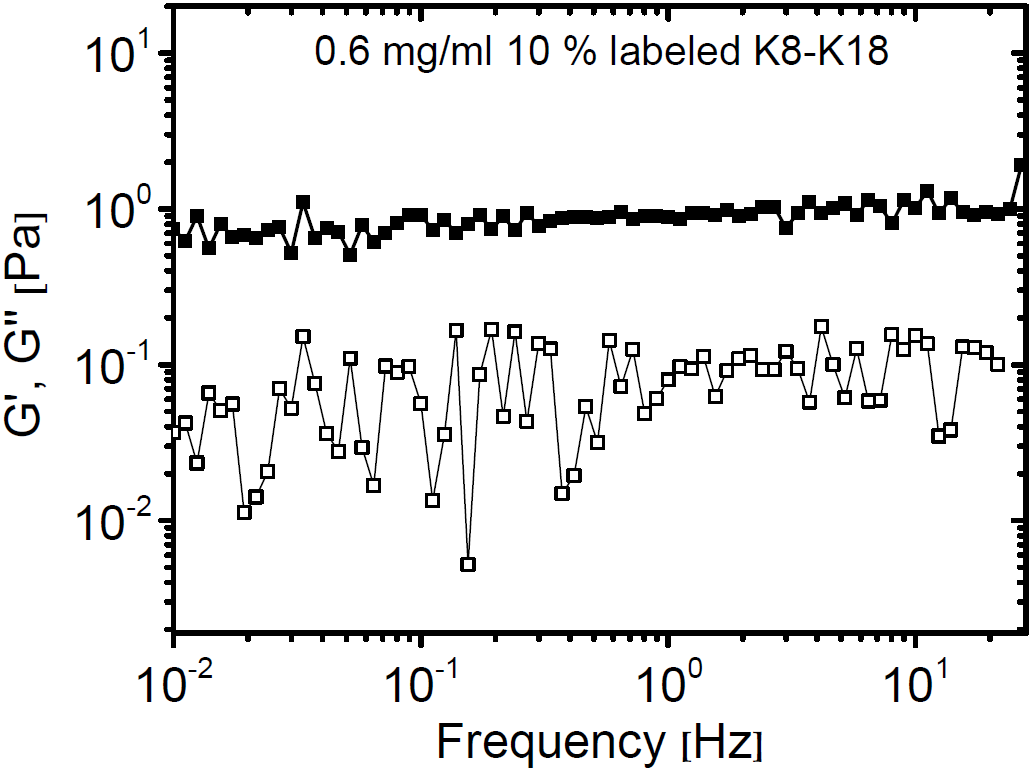 | 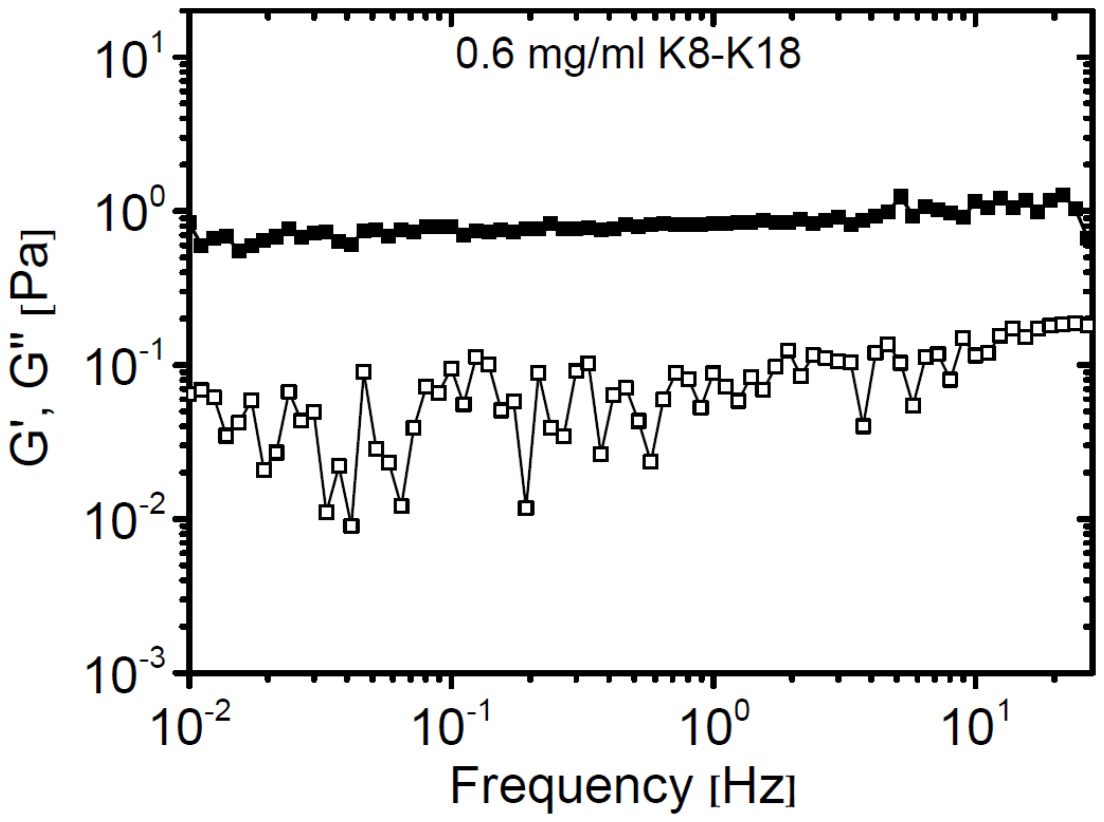 |
| --- | --- |

**Fig. S2**. Linear viscoelastic properties of 10% labeled keratin (0.6 mg/ml) assembled in F-buffer. The labeled fraction did not affect the filaments and network properties. Data points are the mean of 3 independent measurements

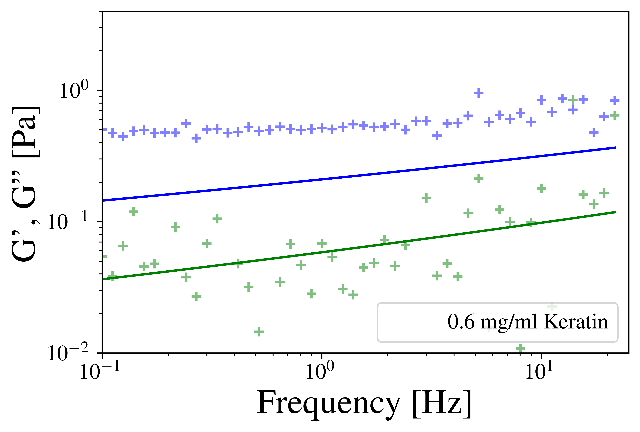

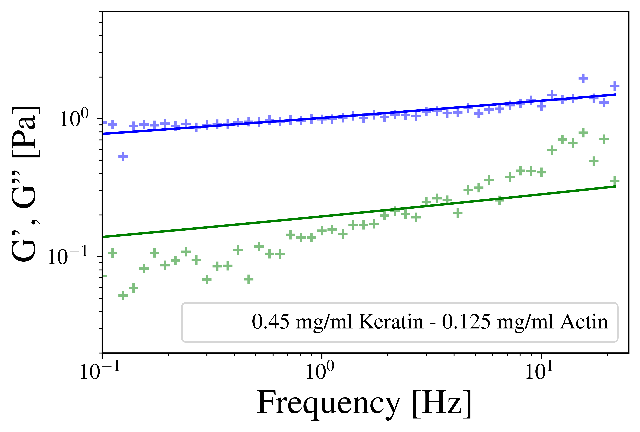

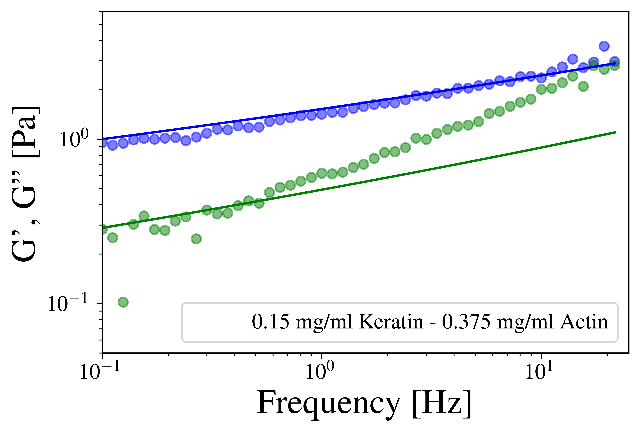

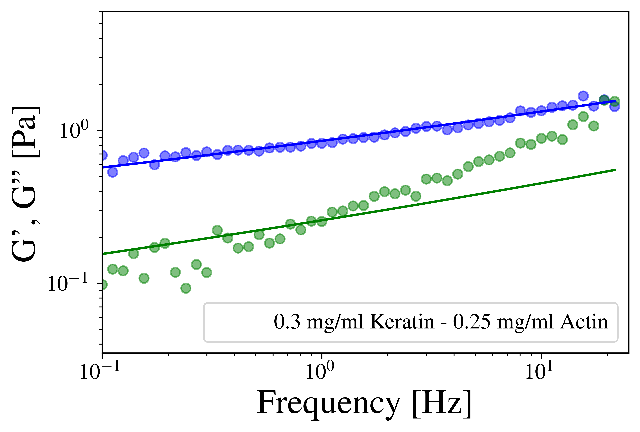

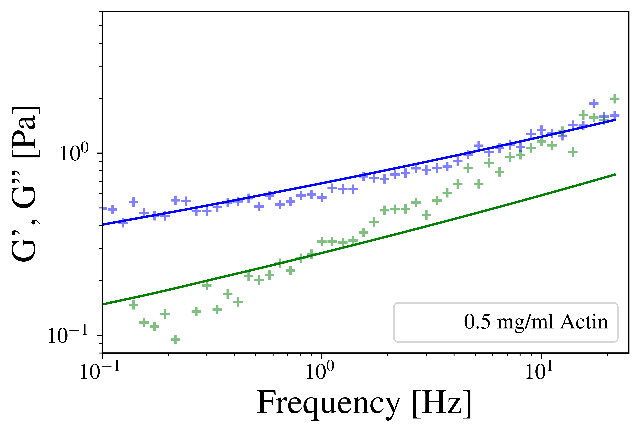

**Fig. S3.** Means of elastic modulus (blue crosses) and loss modulus (green crosses) included GWLC model fits for Keratin-Actin composite networks in compositions of 0.6 mg/ml Keratin (top left), 0.25 mg/ml Actin (middle left), 0.15mg/ml Keratin-0.375 mg/ml Actin (middle right), 0.5 mg/ml Actin (lower left). Model fit parameters for the composites as well as the pure networks are shown in figure 3A and discussed in the main article.

**Significance levels for loss factors:**

### Table S1. P-values obtained from a Mann–Whitney *U* test

|  | 0.6 mg/ml Keratin | 0.45 mg/ml Keratin - 0.125 mg/ml Actin | 0.3 mg/ml Keratin - 0.25 mg/ml Actin | 0.15 mg/ml Keratin - 0.375 mg/ml Actin | 0.5 mg/ml Actin |
| --- | --- | --- | --- | --- | --- |
| 0.6 mg/ml Keratin | - | p=0.114 | p=0.002 | p=0.000 | p=0.003 |
| 0.45 mg/ml Keratin - 0.125 mg/ml Actin | p=0.114 | - | p=0.004 | p=0.000 | p=0.005 |
| 0.3 mg/ml Keratin - 0.25 mg/ml Actin | p=0.002 | p=0.004 | - | p=0.012 | p=0.034 |
| 0.15 mg/ml Keratin - 0.375 mg/ml Actin | p=0.000 | p=0.000 | p=0.012 | - | p=0.255 |
| 0.5 mg/ml Actin | p=0.003 | p=0.005 | p=0.034 | p=0.255 | - |

**Parameters used for figure 5:**

**Table S2**. Parameters used for the calculation of the replicated curves for the nonlinear differential shear modulus in figure 5

|  | Difference between bound and unbound state Δx | Characteristic width of free energy well δ |
| --- | --- | --- |
| 0.5 mg/ml Actin | 200 nm | 30 nm |
| 0.15 mg/ml Keratin - 0.375 mg/ml Actin | 200 nm | 1.5 nm |
| 0.3 mg/ml Keratin - 0.25 mg/ml Actin | 20 nm | 1 nm |
| 0.45 mg/ml Keratin - 0.125 mg/ml Actin | 20 nm | 1 nm |
| 0.6 mg/ml Keratin | - | - |

The energy difference between the bound and unbound state U = 2.5 $k_{B}T$, as well as the control parameter for filament lengthening S = 0.0 were the same for all composites.

**SUPPORTING REFERENCES**

1.	T. Golde, C. Huster, M. Glaser, T. Handler, H. Herrmann, J. A. Kas and J. Schnauss, *Soft Matter*, 2018, **14**, 7970-7978.

2.	K. Kroy and J. Glaser, *New. J. Phys.*, 2007, **9**, 416-416.

3.	T. Golde, M. Glaser, C. Tutmarc, I. Elbalasy, C. Huster, G. Busteros, D. M. Smith, H. Herrmann, J. A. Käs and J. Schnauß, *Soft Matter*, 2019, **15**, 4865-4872.
